# Early versus late noise differentially enhances or degrades context-dependent choice

**DOI:** 10.1101/2024.03.26.586597

**Authors:** Bo Shen, Duc Nguyen, Jailyn Wilson, Paul W. Glimcher, Kenway Louie

## Abstract

Noise is a fundamental problem for information processing in neural systems. In decision-making, noise is thought to cause stochastic errors in choice. However, little is known about how noise arising from different sources may contribute differently to value coding and choice behaviors. Here, we examine how noise arising early versus late in the decision process differentially impacts context-dependent choice behavior. We find in model simulations that under early noise, contextual information enhances choice accuracy, while under late noise, context degrades choice accuracy. Furthermore, we verify these opposing predictions in experimental human choice behavior. Manipulating early and late noise – by inducing uncertainty in option values and controlling time pressure – produces dissociable positive and negative context effects. These findings reconcile controversial experimental findings in the literature, suggesting a unified mechanism for context-dependent choice. More broadly, these findings highlight how different sources of noise can interact with neural computations to differentially modulate behavior.

## Introduction

Classical rational decision theory requires that choices and choice accuracy should be context-independent. For instance, a preference for an apple over an orange should not be influenced by the presence of a banana - a property known as *independence of irrelevant alternatives*, or *IIA*^1,2^. Nevertheless, a wealth of empirical evidence gathered in species ranging from humans to insects has shown that real-world decision-making is inherently context-dependent. The presence of an additional option, even when this option is never chosen, will change choice behavior between other options in the choice set^3–13^. Context-dependent choices are widely documented in choice tasks involving multiple attributes, where the options differ in multiple feature dimensions^14,15^. Beyond that, recent findings show context-dependent preferences can arise even when option values are unidimensional^9,16,17^. Such value-driven effects suggest that contextual processing is a general feature of biological decision-making, but the specific neural mechanisms behind context-dependent choice remain unclear.

One promising explanation for context-dependent choice is the divisive normalization computation, a widely documented nonlinear neural computation seen in diverse species, brain regions, and cognitive processes, that has been proven to be optimal for many environments^18^. The ubiquity of this computation suggests it is canonical and pervasive in neural systems^9,17,19–22^. Functionally, divisive normalization adjusts the representation of input-driven activity in a neuronal pool by dividing it by the activity from other inputs, a general feature required of all optimal systems that have limited capacity^23^. This operation leads to a form of contextual suppression. Additional contextual inputs inhibit the activities driven by the pre-existing inputs and can thus lead to reduced precision in the internal representation. In sensory brain areas, normalization-driven contextual suppression leads to phenomena like surround suppression^19,24^; in decision-related brain areas, normalization produces relative value coding^17,25,26^. In choice, contextual suppression reduces the neural representation of option values when additional contextual options are present. This suppression predicts that choice accuracy between two fixed options will decrease as a function of the value of a third distracter alternative in the choice set. This has been referred to as a negative distracter effect and has been observed experimentally in animals and humans^4,7,9,13,16,27,28^. Analogous extended normalization models explain multi-attribute context effects^29–31^, suggesting that divisive normalization plays a central role in context-dependent preferences.

However, recent empirical studies have also begun to document contextual effects on choice accuracy at odds with the predictions of the standard normalization model. Some studies have reported no overall effect of distracter value on choice accuracy^32^, while other studies have identified a positive distracter effect: at least in certain conditions, the value of a contextual option can enhance the accuracy of choosing between other options^4,5,33,34^. Empirically, both positive and negative distracter effects can coexist in the same experiment, leading to the proposal that the two phenomena reflect different circuit mechanisms, perhaps operating in different brain areas^35^ and/or driven by different individual traits^36^. These observations challenge the standard normalization-based understanding of context-dependent preferences and underscore the need for a unifying framework for all context effects.

Here, we extend the standard framework, adding a richer definition of neural noise. We find that with this richer definition of noise, a richer set of context effects should be observable. Standard normalization models of stochastic choice behavior represent noise as being added to the internal representation after the normalization process is complete, implying that the value inputs into the normalization process must be noiseless and thus deterministic. However, this assumption overlooks the fact that neural value representation is inherently uncertain, with uncertainty inherited from a noisy sensory environment^37^ or originating from internal processes such as inference or memory retrieval of reward-related information^38–40^. It has been previously noted that noise arising early or late in the decision-making process can lead to distinct impacts on choice under specific neural computations^41^. Here, we extend that notion, showing that the stage at which noise enters the valuation process — early in the process of representing values prior to normalization or later during the comparison process post-normalization — can critically influence the direction of observed context effects.

We find theoretically that a context-setting option (for example a low-valued distracter) can actually improve choice accuracy by suppressing early noise. This stands in contrast to the detrimental effect of the same context-setting option in the presence of more traditional post-normalization noise. To empirically validate our predictions, we devised an experimental design that allowed us to tease apart the impacts of early and late noise on the contextual effects of decision-making. Our theoretical and empirical findings reveal that when early noise dominates, a contextual item enhances choice accuracy; in contrast, when late noise dominates, a contextual item impairs choice accuracy. These results reconcile a broad and conflicting literature^4,5,9,32,34^, supporting a canonical role for normalization in neural computation and emphasizing the critical importance of different types of uncertainty in neural coding.

## Results

### Divisive normalization under different noise sources

While it is known that divisive normalization shapes the noise structure of neural response variability^42–44^, how neural noise and normalization work together to affect cognitive processes and behavior is unknown. Here, we examine how neural noise arising from different stages of value normalization affects context effects in decision-making: *early noise*, which arises prior to the normalization process (**Figure 1a)**, and *late noise*, which arises after normalization (**Figure 1e)**. We conceptualize early noise as the variability inherent in the input values feeding into the normalization process. This variability captures uncertainties in a subject’s ability to assign a valuation to an individual item, which can result from the imprecision of sensory inputs, the vagueness of reward-related associations, and the fluctuation of the decision-maker’s motivational states. In contrast, we conceptualize late noise as the stochasticity of neural processing after normalization, for example, in post-normalization value coding or when performing a choice comparison amongst multiple options that have already been efficiently represented. Such late noise processes have been a longstanding component of both economic stochastic choice theories^45–48^ and neurocomputational models of decision-making^49–51^. At the neural level, late noise is typically considered to represent variability in the firing rates of value-coding neurons, independent from the inputs and participating in the winner-take-all selection process^52,53^.

**Figure 1.**
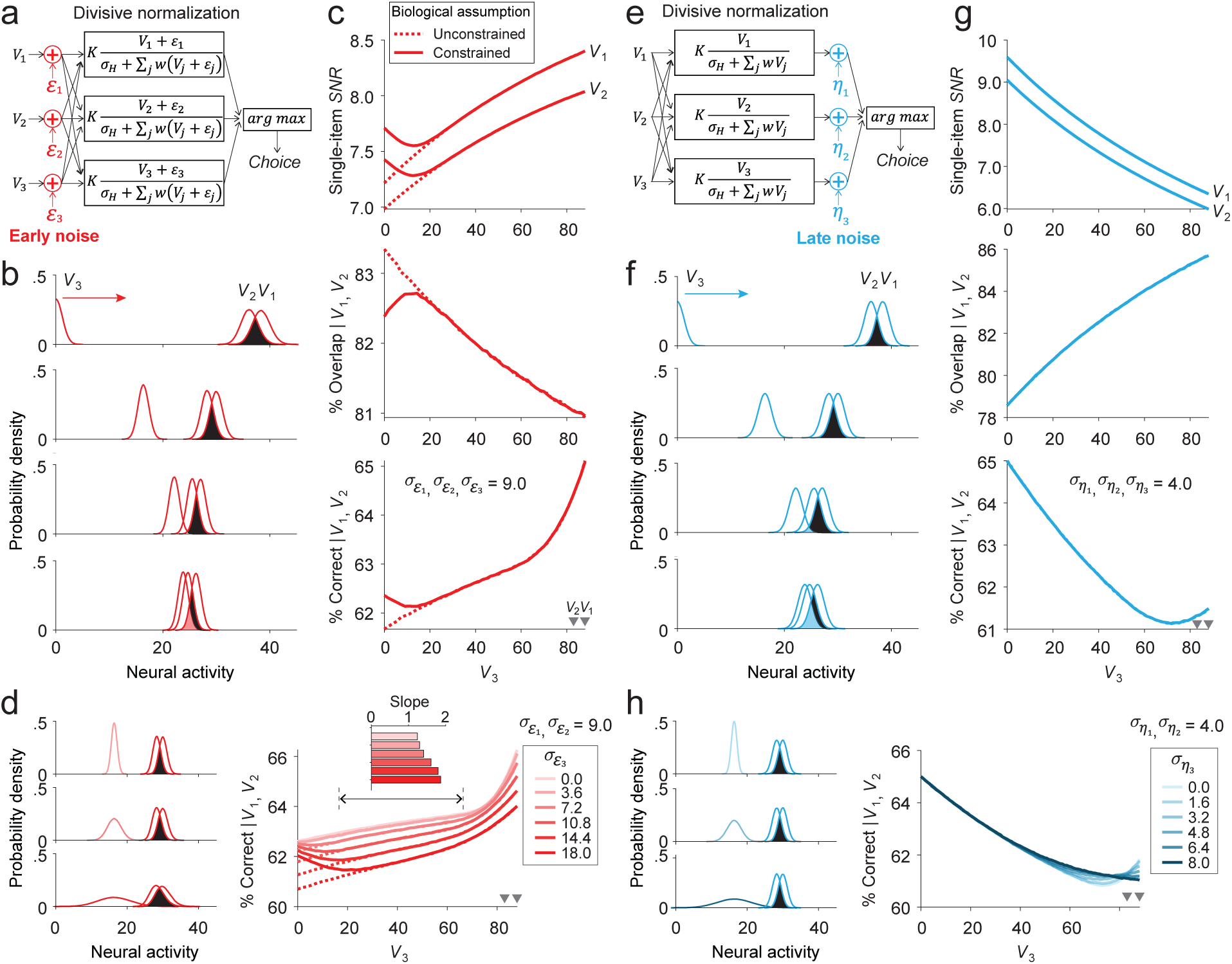
Divisive normalization leads to opposite contextual effects under early and late noise. **a** An illustration of the divisive normalization model with early noise. **b** With only early noise, increased contextual value induced by a distracter (*V_3_*) reduces the mean activities of the targets (*V_1_* and *V_2_*) by divisive normalization, meanwhile sharpening the target activity distributions. **c** The signal-to-noise ratio (*SNR*, measured as the ratio of mean over standard deviation) of every single target activity is improved by *V_3_* when under early noise (upper panel); the percentage of overlap between the neural activities of the two target distributions decreases (middle panel). Thus, the divisive normalization model under early noise predicts a positive context effect where *V_3_* facilitates the conditional choice accuracy between the targets (bottom panel). A non-negative threshold was applied for the biologically constrained simulations (solid curves), whereas activities were allowed to be negative in the unconstrained assumption (dashed curves). A steeper increase on the right end of the x-axis was observed due to competition between the distracter and the lower-value target (see text). **d** With only early noise, fixing the mean and variance of the targets and testing the contextual effects at multiple levels of the distracter’s variance uncovers an impairment effect of contextual variance on the target conditional target choice accuracy (graded colors in the right panel). However, nosier distracters paradoxically demonstrate steeper positive contextual effects. Slopes of the curves visualized in the inset were quantified between the middle range of *V_3_* (scaling between 0.2 and 0.8 relative to *V_2_*) to roughly get rid of the impacts of the biological cut on the left and the competition effect on the right. **e** An illustration of the divisive normalization model with late noise. **f** With only late noise, increased *V_3_* reduces the mean activities of the targets but has no impact on the sharpness of the target activity distributions. **g** The *SNR* of the target activities is impaired (upper panel), and the percentage of overlap between the targets increases (middle panel), thus leading to a negative context effect where *V_3_* impairs target choice accuracy under late noise (bottom panel). **h** Testing multiple levels of the distracter’s late noise on fixed targets does not lead to an impact of contextual variance shown under early noise. The curves overlap and consistently show negative context effects. Under both early and late noise, the choice accuracy curves exhibit an increasing ‘hook’ when *V_3_* approaches *V_1_* and *V_2_* due to a different non-normalization mechanism (see text).

To theoretically examine the differential impact of early and late noise in context effects, we extended a previous algorithmic model of divisive normalization for stochastic choice behavior^9^. In this model, the represented value of each option (*FR_i_*) is conceptualized as neural activity, akin to the firing rates of a neuronal pool, driven under the direct input of the option (*V_i_*) and normalized by the integration of the input from all options (Equation 1). Unlike the previous model, which assumes no noise on the input values (*V_i_*), we included an early noise term regulating the variability of each input. This early noise is defined by a zero-mean *Gaussian* distribution additive to its mean value, 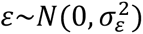. Late noise is then added onto the product of divisive normalization as drawn from another zero-mean *Gaussian* distribution, 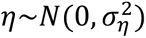, independent from early noise. A decision is subsequently made by choosing the option with the largest represented value under the two sources of noise; see *Methods for* other details of implementation.

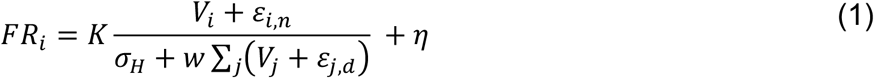

where *K* is the maximum neural response, *σ*_*H*_ is a semi-saturation constant determining the baseline normalization, *w* is the weight of the normalization. *ε*_*i*,*n*_ and *ε*_*i*,*d*_ are the early noise components of the option *i* when in the numerator and the denominator, respectively. They share the same distribution but were marked differently since the covariance between them will moderate the early-noise-driven effect proposed in the following text (see analytical analysis of covariance in *Supplementary Note 1: Under early noise*). Considering that the denominator-driven normalization likely arises from biophysical or circuit mechanisms independent of feedforward input, resulting in independent noise between the numerator and the denominator^24,54–56^, we applied independent noise in our following analyses. *η* is the late noise, which was assumed to be independent of early noise since the two terms reflect variation from two independent sources. Across options, we assumed both early noise and late noise are independent. Covariance of noise across options can change the stochasticity of choice behavior, which has been well documented in the literature and is not covered under the scope of the current analyses^47,57^.

### Normalization under early and late noise predicts opposite contextual effects

To examine how early and late noise contribute to context-dependent choice, we simulated the classic trinary choice decision-making experiment of Louie and colleagues^9,32^. In this widely used approach to quantifying contextual choice effects, two higher-value options (*V_1_* and *V_2_*), *the targets*, are used to probe the impact of a context-setting option (*V_3_*), *the distracter*. In this framework, target-coding neural activities consist of two distributions, and stochastic choice between targets arises from the overlap between those distributions. Context effects are evident when the value of the distracter option changes the overlap between target distributions (see visualization examples in **Figures 1b and 1f**).

In previous work, stochastic choice and the contextual effect of the distracter are presumed to derive from late noise^5,9,13^. With such post-normalization late noise, a contextual input suppresses the mean activities of other options in the choice set, leading to diminished discriminability between target activities and, consequently, impaired choice accuracy. Larger distracter values lead to more impaired choice, therefore producing a negative context effect, which has been shown in previous computational and empirical data^6,9,13^. Here, we reproduced the original visualization of this process in **Figure 1f**. This reduction in choice accuracy is evident in the signal-to-noise ratio (*SNR*, measured as the inverse of the Coefficient of Variance) of individual option representations: when the distracter value increases, the model representation of each target becomes relatively noisier (**Figure 1g**, upper panel). Further, given the divisive nature of normalization, increasing contextual value reduces the space between the two target distributions and, therefore, enlarges their amount of overlap (**Figure 1g**, middle panel). At the behavioral level, these changes lead to a decrease in conditional choice accuracy (relative choice accuracy between the two target options, excluding distracter choices) (**Figure 1g**, bottom panel). Note that when the distracter value increases to the point at which it becomes roughly comparable to that of the targets, a ’hook’ in the accuracy curve occurs because the distracter begins to compete selectively with the second-best target; decreasing the fraction of trials on which participants choose the second-best target naturally increases the relative ratio of choosing the best target over the second-best target. This phenomenon of *conditional choice accuracy* has been well explained in previous research^9^ and is not specific to divisive normalization.

In contrast to these well-described effects, we report here that when noise is introduced prior to normalization, distracter values can enhance choice accuracy. Under early noise, increasing the mean value of a contextual input (*V_3_*) while keeping other variables fixed improved the *SNR* of the normalization denominator (*σ*_*H*_ + *w*(*V*_1_ + *ε*_1_ + *V*_2_ + *ε*_2_ + *V*_3_ + *ε*_3_)). This improvement effectively leads to an improved representation of individual targets against the suppression of the target’s mean, evident as sharpened activity distributions (**Figure 1b**) and increasing *SNR* of each individual target (**Figure 1c**, upper panel). The improved representation leads to decreased overlap between the target distributions even when the distance between their mean values is reduced (**Figure 1c**, middle panel). Overall, this effect improves conditional choice accuracy (**Figure 1c**, bottom panel). Thus, the interaction of early noise and normalization predicts an increase in choice accuracy with larger distracter values - a positive distracter effect.

Beyond the simulations, our analytical analysis showed that this effect is general (see *Supplementary Note 1: Under early noise*). Increasing the distracter’s mean value improves the target’s mean value relative to its variance as a distribution of the *Ratio of Gaussians*, where both the numerator and the denominator in Equation 1 are *Gaussian* distributed^43,58^. The impact of *V_3_* over the discriminability of the targets stays positive when the choice only involves early noise (occurring on any options) and the target values are above zero (see proof in *Supplementary Note 1: Under early noise*). However, it is important to note that increasing the early noise of the distracter will always decrease the choice accuracy between the targets, evident from the decreased accuracy across the graded lines in **Figure 1d** (see proof in *Supplementary Note 1: Under early noise*). However, the positive contextual slopes are paradoxically steeper when under noisier distracters, since increasing the mean value of the distracter contributes more to the *SNR* of the normalization denominator. In contrast, when the distracter’s variance occurs in late noise, the contextual variance does not contribute to contextual modulation, evident from the overlapped curves in **Figure 1h** (see proof in *Supplementary Note 1: Under early noise*), signifying the fundamental difference between early and late noise. In summary, the positive effect we showed here under early noise is due to the increase of the mean distracter value relative to the overall early-noise variance in the normalization denominator, which improves the representation and discriminability of the targets. Furthermore, it is important to assume independence between the noise terms, across options and within options. For example, assuming positive within-option covariance between the numerator and the denominator of Equation 1 would compromise the positive effect (see details in *Supplementary Note 1: Under early noise*; and see the simulation details in *Methods*).

When applying a (biological) constraint that all values below zero must be represented as zero, i.e., the *non-negative constraint for neural activities*, we observed subtle differences in model predictions (solid lines in **Figure 1c**) compared to the implementation allowing negative neural activities (dashed lines). Since distracter values are lower than target values by design, imposing a non-negative constraint primarily reduces the variance of the distracter option. This, in turn, improves target representation *SNR* and increases conditional choice accuracy when the distracter value approaches zero. Such an effect exclusively arises under early noise; applying non-negative constraints under late noise does not appreciably change the effects of context on representations or choice (**Figure 1g**). Although it is not the main focus of the current study, the impact of non-negative constraints on contextual processing highlights the impact of biological constraints during the computational operations of neural noise^44,57,59^.

### Divisive normalization with mixed noise can replicate all context effects

We focused above on the distinct effects of early and late noise separately, whereas, in the brain, both types of variability likely co-exist. We thus examined predictions under mixed noise by integrating both sources of noise into the normalization model. We found that opposing contextual effects compete and progressively change from improving to impairing choice accuracy when early and late noise tradeoffs in the model (**Figure 2a**). When early noise predominates, distracter value facilitates choice, evident from the upward trend of the curves (red-to-orange). Conversely, increasing late noise progressively shifts the context effect to impairing choice accuracy, as indicated in the downward slope of the curves (green-to-blue). Our analytical approach confirmed such competition by showing that the overall degrees of early noise across all options compete with the summed late noise of the targets in driving positive and negative context effects, respectively (see *Supplementary Note 1: Under mixed noise*). Therefore, a divisive normalization model with two sources of noise can replicate the range of context effects reported in the literature, either positive, negative, or non-effects^4,5,9,32,34,35^.

**Figure 2.**
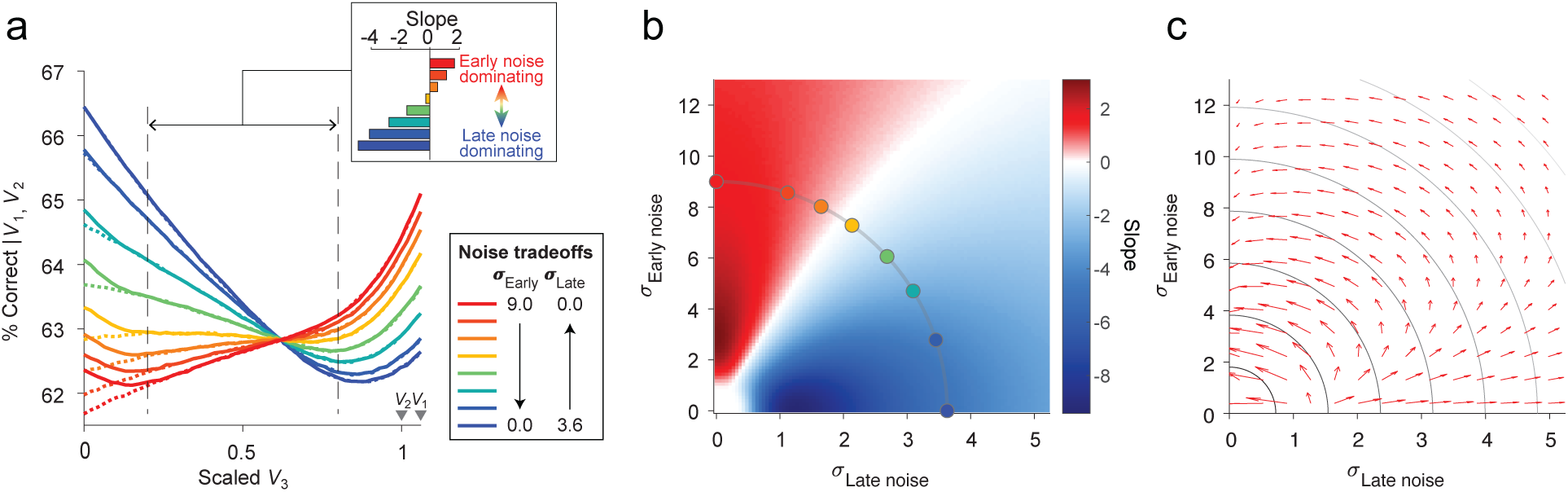
Context effects under mixed noise. **a** Early and late noise compete in driving opposite trends of context effects. In the simulations, early noise was reduced from red to blue curves; meanwhile, late noise was increased from blue to red curves. The degrees of early and late noise were set as equal for every option. The inset illustrates the regression slope between the middle range of scaled *V_3_*, from 0.2 to 0.8, to eliminate the impacts when the distracter value approaches zero or the value of the targets. **b** Early and late noise tradeoffs in a wide range of noise magnitudes. The quantified slopes from the middle range of *V_3_*. Spots along the line have a constant Euclidean norm of the two sources of noise, which were the set parameters visualized in panel **a**. **c** The vector field indicates the changing trend of context effects at every combination of early and late noise. The vectors consistently point towards the directions where early noise increases and late noise decreases, along the lines of constant Euclidean norm (lines in grayscale).

To quantify the context effects that were consistently different between early and late noise conditions, we reasoned that analyses should exclude the ranges of contextual values near zero or near the target values, where simulations show complicated curvatures that are non-specific to normalization and similar effects under both early and late noise. We thus focused on quantifying context effects as the slope of the curves in the middle range of *V_3_* (between .2 and .8 when *V_3_* is scaled to the minimum value of the targets), where both impacts of biological cutoff and distracter-target competition effect are small (< 3.3% for the left end biological cutoff and < 7.5% for the right end competition effect in our case, but the percentages would vary when the options’ variance changes). This measure shows a clear pattern of progressive change in context effects as the degree of noise shifts from early-dominant to late-dominant (inset bar graph, **Figure 2a**).

How robust is the tradeoff between the predicted context effects when early and late noise compete? Using the same quantification approach as above, we visualized predicted context effects across a wide range of early and late noise magnitudes (**Figure 2b**). These results support the general conclusion that larger early noise drives positive context effects (red) while larger late noise drives negative context effects (blue) (see proof in *Supplementary Note 1: Under mixed noise*). In addition to this general pattern, ceiling and floor effects occur when both sources of noise are small (bottom-left corner) or large (upper-right corner); in these regions, accuracy is either very low (near random choice) or saturates to 100%, minimizing the potential to observe context effects. By controlling noise magnitudes, we find a clear pattern of tradeoffs between early and late noise effects along the lines of a constant Euclidean norm (lines in **Figure 2c**) of the two sources of noise. In other words, given a constant sum of the two sources of noise, decreasing late noise and increasing early noise always increases the context effect, and vice versa. Thus, the observed variability in contextual effects, contingent on the balance of early and late noise, offers a testable empirical approach to validate our proposed framework.

### Model-driven experimental design

To test the predictions of our model, we designed an experiment to systematically manipulate and quantify the influences of early and late noise on context effects. A 2-by-2 factorial design was employed to create four conditions in a within-subject design that combined varying degrees of early and late noise (**Figure 3a**). We adapted a two-stage task used in previous studies^9,32^. In our version of this task, participants (*N* = 55) were first presented with different consumer items (e.g., drone, tea kettle, or camera; **Figure 3b**) and provided their subjective monetary valuation for each item (**Figure 3c**) in an incentive-compatible procedure. Subsequently, participants were presented with trinary choices pairing two high-valued items and a distracter of varying value (**Figure 3d**). They were asked to choose their preferred item from these sets of three items (**Figure 3e**).

**Figure 3.**
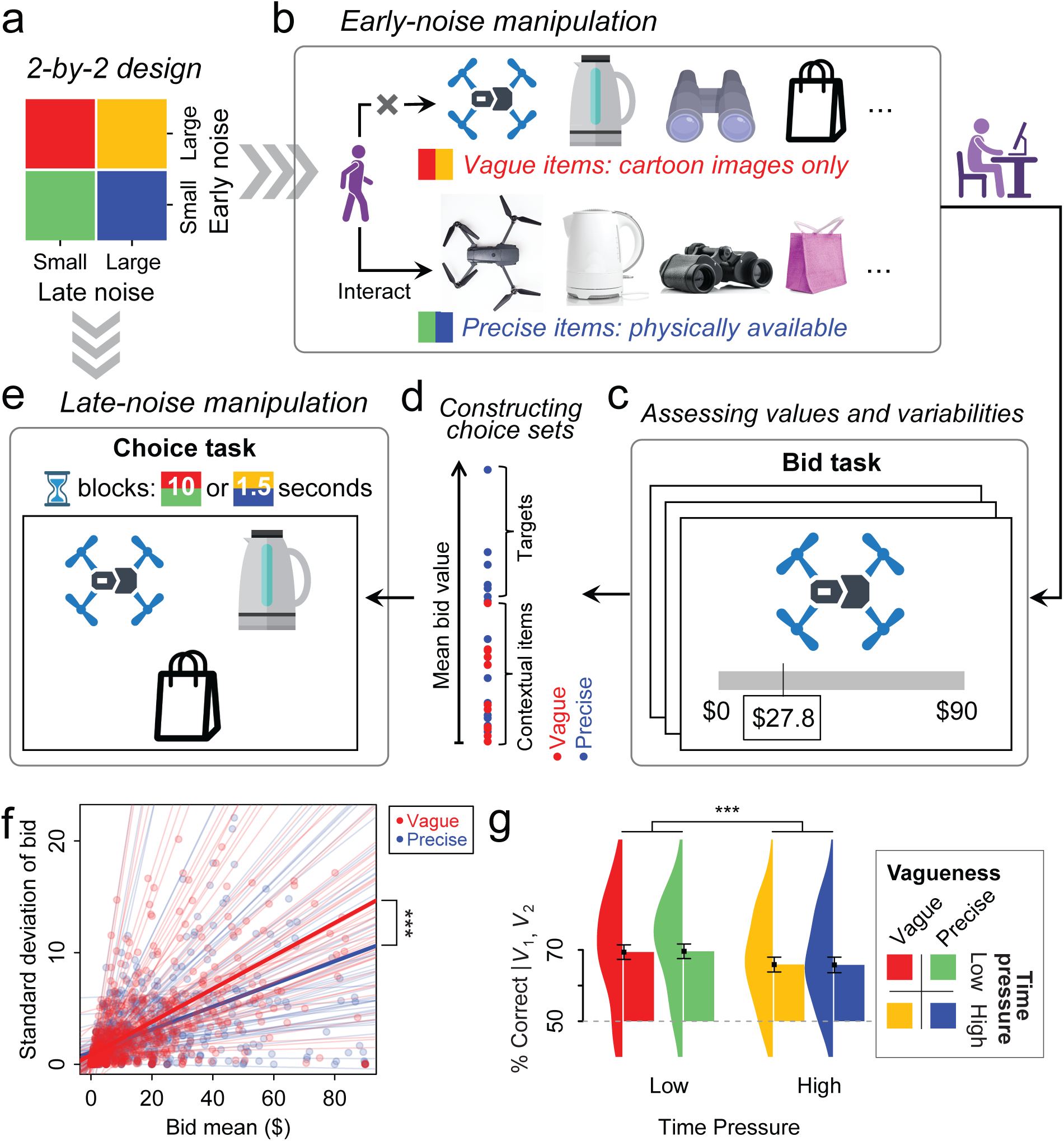
Experimental design and procedure in testing context effects under different sources of noise. **a** A 2-by-2 design matrix was used to independently vary the levels of early and late noise. **b** Early noise was manipulated by presenting the consumer goods either physically (handled and viewed) or only as cartoon images. This induced two different degrees of precision in object valuations. **c** To reveal the subjective value and the noise associated with the representation of each item, participants were asked to BDM-bid on each item three times. **d** To build the choice sets for the subsequent task, a fixed set of the targets (*V_1_* and *V_2_*) was constructed from the six highest-bid precisely presented items, resulting in 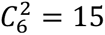 combinations of target pairs. In each trial, one target pair was combined with one distracter (*V_3_*) selected from twenty-four lower-bid items, where half of them were precisely presented items. **e** In the choice task, the participants were asked to choose their preferred item from a triplet under low-(10 seconds) and high-(1.5 seconds) time-pressure blocks. All images were presented as cartoons, even when they were precise items to control for visual confounds. The positions of the options were randomized across trials. **f** The effect of the manipulation on early noise confirmed that the standard deviation of bidding increased faster with the bid mean in the vague items than in the precise items (*\*\*\**, 55 participants, linear mixed regression: *β* = .049, s.e. = .011, *t* = 4.52, *p* < .001, standardized coefficient = .13, 95% C.I. = [.05, .20]). Each dot indicates an item; thin lines indicate fitting to each individual, and thick lines indicate group averages. **g** The effect of the manipulation on late noise confirmed that the participants’ choice accuracy decreased when time pressure increased (***, 55 participants, within-subject ANOVA: *F*_1, 162_ = 39.17; *p* < .001, partial *η*^2^ = .19, 95% C.I. = [.11, 1.00]). Bar heights and whiskers represent the mean and s.e. of choice accuracy; the distribution is attached on the left of each bar. (Human-shaped and hourglass icons are created in BioRender. Shen, B. (2025) https://BioRender.com/s83i567.)

The degree of early noise was modulated by presenting items to participants with varying levels of vagueness. To elicit valuations that were as precise as possible (and thus to reduce early noise), half of the consumer items were physically present in the laboratory and the participants were required to handle them. We hoped in this way to reduce early noise by providing enough information for participants to form precise evaluations of their desirability. To elicit variable valuations (and thus to increase early noise), the other half of the consumer items were not physically present and were depicted only as cartoon images. This had the effect of rendering their specific characteristics and quality uncertain (**Figure 3b**). Following interaction with the items, participants’ valuations were assessed through a Becker-DeGroot-Marschak (BDM) auction task, as depicted in **Figure 3c**. Participants were asked to bid three times for each item, all of which were visualized as cartoons to prevent visual confounds. This allowed us to characterize the variability of valuations as a proxy for early noise (see *Methods* for details). Bids for all items demonstrated a variance that depended on the mean bid. As expected, the variance-mean relationship was steeper for the vaguely described items than for the items handled by the participants (difference of slopes: *β* = .049, s.e. = .011, *t* = 4.52, *p* < .001, standardized coefficient = .13, 95% C.I. = [.05, .20]; **Figure 3f**). This affirmed the effectiveness of our early-noise manipulation. The mean bids between the vague and precise conditions showed no significant difference (Repeated-measurement ANOVA, *F*_1,1924_ = 2.15, *p* = .143, partial *η*^2^ = 1.12×10^-3^, 95% C.I. = [0, 1.00]).

The three-item choice sets were constructed based on the early noise structure for our context experiment (**Figure 3d**). To test whether the context effect under early noise matches our prediction, we varied the mean and the early noise of the distracters and kept a fixed set of targets. In this way, any change of choices between the targets will be contributed by the distracter – a well-controlled situation to focus on contextual modulation. Thus, we picked distracter items having a range of values, but sorted into those handled by the participants (precise) and those represented only by cartoons (vague). The distracters showed a steeper variance-mean slope in the vague condition than the precise condition (*β* = .034, s.e. = .011, *t* = 3.18, *p* = .001, standardized coefficient = .13, 95% C.I. = [.05, .20]) but no significant difference in their mean bids (*F*_1,1264_ = .039, *p* = .84, *η*^2^ = 3.1×10^-5^, 95% C.I. = [0, 1.00]) or chosen ratios (*F*_1,54_ = .58, *p* = .45, *η*^2^ = .01, 95% C.I. = [0, 1.00]) between the conditions, affirming that they were well-manipulated and controlled. In contrast, all target options (the higher-valued pair in our design) were picked as a fixed set of items that had been seen and handled by the participants (members of the precise pool). All options were presented as cartoons even when they were precise items to control for visual confounds.

To manipulate the degree of late or decisional noise, we imposed a decisional time pressure, requiring the participants to make their choices either quickly or slowly (in a block-wise fashion). This is an approach widely used to increase decisional stochasticity^60^. In the low time-pressure block, the participants had 10 s to make a decision (participants took 1.17 ± .05 s, with .06 ± .04% time out); in the high time-pressure block, the participants had only 1.5 s to choose (participants took .78 ± .02 s, with 3.72 ± .66% time out; responded significantly faster than under low-pressure, *t*_54_ = -9.91, *p* < .001, Cohen’s D = -1.24, 95% C.I. = [-1.59, -.88]). This is indicated by a color code in **Figure 3e**. Consistent with an effect on late noise, time pressure significantly reduced participants’ overall decision accuracy across conditions (*F*_1, 162_ = 39.17; *p* < .001, partial *η*^2^ = .19, 95% C.I. = [.11, 1.00]; **Figure 3g**).

### Model comparison of empirical data reveals divisive normalization with two stages of noise

Using the design above, we examined the effects of early and late noise on contextual choice behavior by comparing predictions across four alternative models. These models ranged from a basic probit choice model with linear, independent value coding, to more complex models incorporating divisive normalization and two stages of noise (see model specification in *Methods*). In simulations, we fixed the targets and varied the mean values of the distracter with multiple levels of variance to manipulate their early noise. Two levels of late noise were applied to all options, matching the time pressure design. The simplest probit model (*Model 1*) showed only the effect of late noise on the conditional choice accuracy between the targets (as an overall vertical shift in accuracy), but no contextual modulation effect as *V_3_* increases (**Figure 4a**). Including early noise (*Model 2*; **Figure 4b**) yielded the same flat prediction for contextual modulation; however, earlier emergence of the “hook” under higher *V_3_* noise resulted from the choice competition between the distracter and the lower-value target, a different mechanism from contextual modulation described above. When value coding was adapted to divisive normalization but included only late noise (*Model 3*, the classical divisive normalization model), it predicted negative contextual modulation, with late noise affecting overall choice accuracy but no impact from early noise (**Figure 4c**). *Model 4*, which applied divisive normalization with both early and late noise (**Figure 4d**), uniquely predicted an effect of contextual early noise on target choice. As early noise in *V_3_* increased, target choice accuracy decreased, marking a unique feature of contextual modulation where both contextual mean and contextual variance play a role. Meanwhile, the contextual modulation slopes shifted from negative to positive with increased contextual variance (colored lines within each panel in **Figure 4d**), reflecting the mechanism of contextual facilitation under high early noise as predicted by our theory. Adjusting late noise levels reduced overall accuracy and moderately shifted contextual modulation slopes to negative trends (comparing between the panels in **Figure 4d**), illustrating an early-late noise tradeoff driving these opposing trends in contextual modulation. We note that in the empirical data the degree of early noise appears to scale with mean value (**Figure 3f**), different from the constant levels of early noise tested above. However, versions of *Model 4* that incorporate mean-scaled early noise still predict positive contextual effects and gradually transitions to negative effects as late noise increases (**Supplementary Figure 1**).

**Figure 4.**
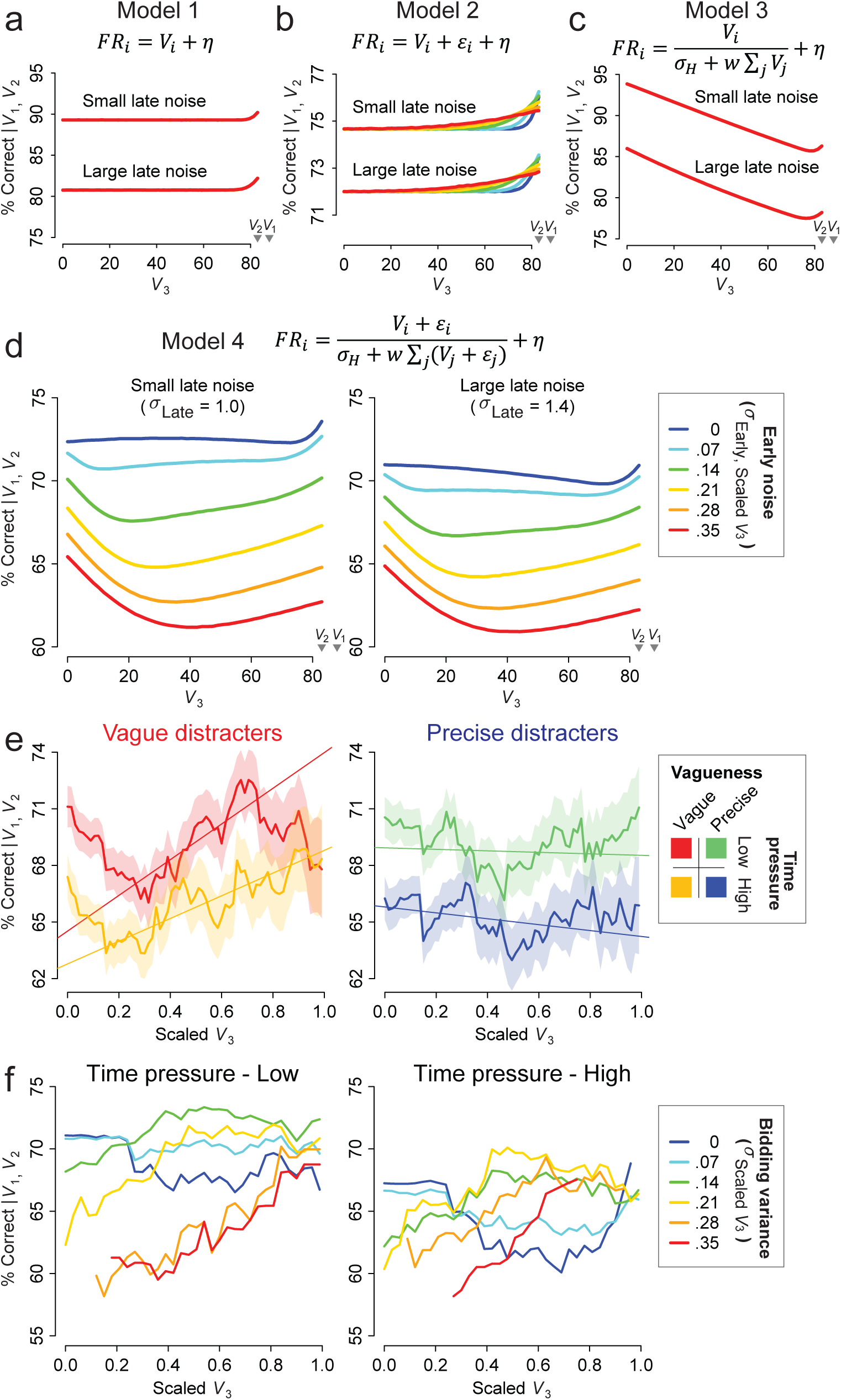
Model comparison of participants’ choices reveals divisive normalization with early and late noise. **a-d** Predictions from different alternative models under the specific experimental design, with fixed pairs of targets and varying distracter mean values (x-axis) and distracter variance (color-coded lines). The early noise of *V_1_* and *V_2_* were fixed to match the overall choice accuracy observed from the data. Two levels of late noise were applied, annotated by text. **e** Context effects on conditional choice accuracy between the targets, aggregated by experimental conditions. All option values were scaled to the minimum target within each subject. Changing trends along the x-axis were disclosed by implementing sliding windows over *V_3_* (window span = .3, step size = .015). The solid lines and the shaded areas indicate the mean choice accuracy and the standard deviation at each sliding window. Linear trends were indicated by the straight lines fitted in the constrained range of scaled *V_3_* (.2 to .8). **f** Sliding windows of choice accuracy across different levels of standard deviation of the scaled *V_3_*, coded in color lines. Each line represents a subset of data within a narrow range of distracter variance (standard deviation ±.07) around the value indicated in the legend, thus a wider window span (.5) and coarser step size (.03) were applied for visualization.

We fitted and compared the four models to participants’ trinary choices. Models were designed in a nested manner to improve the validity of comparison (*Methods*). *Model 2* added a parameter to *Model 1* to incorporate the item-wise early noise, representing the participant’s bidding variance varying across items. This manipulation significantly improved model fitting, suggesting that early noise influences choice stochasticity in an itemized fashion (*Model 2* vs. *Model 1*: ΔAIC = -483; ΔBIC = -272; Table 1). *Model 3*, which assumed only late noise but incorporated divisive normalization on top of *Model 1*, outperformed *Model 1*, suggesting contextual modulation in participants’ choices (*Model 3* vs. *Model 1*: ΔAIC = -459; ΔBIC = -248; Table 1). *Model 4*, which incorporated both divisive normalization and two-stage noise, outperformed all other models in fitting participants’ choices, providing strong evidence for a mechanism including both divisive normalization and two-stage noise (*Model 4* vs. *Model 2* suggesting divisive normalization addition to two-stage noise: ΔAIC = -392; ΔBIC = -181; *Model 4* vs. *Model 3* suggesting two-stage noise addition to divisive normalization: ΔAIC = -415; ΔBIC = -205; Table 1).

To better compare with the existing literature, which largely emphasizes linear trends in contextual modulation, we provided visualizations of these effects along the *V_3_* axis as post-hoc analysis after the model comparisons. It is important to note that the linear regressions presented below have limited statistical validity in capturing the non-linear effects of contextual modulation, as they do not account for multiple factors such as the non-monotonic trends at the two ends of the *V_3_* axis, the early-noise structure of the targets and distracters, or between-subject variance introduced during data pooling. Thus, these regressions should be considered illustrative rather than hypothesis testing.

Within-subject modulation effects are challenging to discern, due to the limited number of trials within subjects (six distracters with uncontrolled distribution of mean values and variance in each experimental condition). To better visualize these effects, we pooled data across subjects by aligning each individual’s option values to their minimal target value and further applied sliding windows along the scaled *V_3_* (window width = .3; step = .015). The pooled data along the *V_3_* axis revealed non-monotonic patterns of contextual modulation within each condition (**Figure 4e**). Linear regression performed on the trends within a constrained range (.2 to .8 scaled *V_3_*) uncovered an effect of distracter vagueness, demonstrating a positive shifting of the contextual slope (*Regression 1*: interaction between vagueness and the mean of scaled *V_3_*: *β* = .59, s.e. = .28, *t* = 2.09, *p* = .037, standardized coefficient = .1, 95% C.I. = [.01, .20]; see *Methods*). Defining distracter’s vagueness by median splitting *V_3_*’s bidding variance resulted in a pattern consistent with the experimental conditions of vagueness (**Supplementary Figure 2**).

The linear pattern, however, did not fully represent the data structure, as each line included the complex structure of mean-scaled contextual variance (as mentioned above in **Figure 3f**). To further dissect this rich data pattern, we aggregated the data into lines within narrow ranges of contextual variance (window width = .14, step = .07 on the standard deviation of *V_3_*’s bidding) (color-coded lines in **Figure 4f**). Contextual modulation under different levels of contextual variance showed varying slopes (*Regression 2*: interaction between distracter mean and distracter variance on conditional choice accuracy: *β* = 1.70, s.e. = .28, *t* = 6.13, *p* < .001, standardized coefficient = .12, 95% C.I. = [.08, .16]; see *Methods*). *V_3_* impaired choice accuracy under low distracter variance (e.g., cyan line on the left panel and blue line on the right panel) and gradually shifted to improve choice accuracy under higher distracter variance (graded color lines within each panel), consistent with our model predictions (**Figure 4d**). A slight shift in intercept for noisier *V_3_* was observed (some lines with nosier *V_3_* at the right end reached higher than the lines of precise *V_3_* in the data, but not in model simulations), due to a random variation in the empirical data during the pooling process (**Supplementary Figure 3a**), which was well-captured by model fitting (**Supplementary Figures 3c and 3d**); simulations confirmed this shift was not a systematic bias introduced by the pooling process (**Supplementary Figure 3b**).

Choice accuracy declined with both distracter mean (*Regression 2*: *β* = -.18, s.e. = .09, *t* = -1.98, *p* = .047, standardized coefficient = .01, 95% C.I. = [-.04, .07]) and distracter variance (*Regression 2*: *β* = -1.04, s.e. = .23, *t* = -4.57, *p* < .001, standardized coefficient = -.11, 95% C.I. = [-.19, -.04]), consistent with an important model prediction that both contextual mean and contextual variance impact choice behavior. In addition, time pressure reduced overall choice accuracy (*Regression 2*: *β* = -.17, s.e. = .05, *t* = -3.55, *p* < .001, standardized coefficient = -.17, 95% C.I. = [-.23, -.10]). Within a constrained range (scaled *V_3_* < .8), time pressure exhibited a marginally significant interaction with contextual variance (*β* = .46, s.e. = .23, *t* = 1.95, *p* = .051, standardized coefficient = .08, 95% C.I. = [0, .15]) — a pattern consistent with the model prediction of a flatter early noise effect under high time pressure due to early-late noise tradeoff.

Together, examining linear trends by considering both contextual mean and contextual variance allowed us to dissect the complex patterns of contextual modulation, demonstrating how participants employed divisive normalization with both early and late noise in their choices.

## Discussion

Variability is a central feature of sensory information, neural coding, and behavior, but how cognitive noise and neural computation interact to control behavior is unknown. Here, we examined how different types of noise interact with divisive normalization-style models. We found that divisive normalization models that incorporate noise in their inputs can replicate the full range of observed context effects. With early noise arising in the inputs to the normalization process, a contextual option can improve the representation of pre-existing options, thus enhancing choice accuracy. In contrast, noise added to the output of the normalization process can react to a contextual option in a way that impairs the representation and thus reduces choice accuracy. These model predictions were validated in empirical human choice behavior, in a task designed to dissociate the effects of early and late noise. Our behavioral findings illuminate the role of noise on choice and suggest that an enhanced divisive normalization model can account for empirically observed effects of early and late noise.

In contrast to the assumptions of classical normative choice theories, the preferences of human and animal choosers change in response to context. Such violations of traditional rational choice models have been widely studied in the economics and psychology literature and are of increasing interest in neuroscience. However, the scientific dialogue on context-dependent effects has been recently marked by controversy, especially regarding how the presence of a contextual option influences the accuracy of decision-making. The classical divisive normalization model requires that a contextual option impairs choice accuracy by diminishing the magnitude of the neural representation of the target options—a view substantiated by a breadth of studies across various species and cognitive tasks^7,9,11,16,17,34,61,62^. However, more recent work suggests that these context effects can be variable. One study failed to replicate negative context effects, finding no effect of distracter value on choice and arguing that negative effects could arise from confounds in analysis^32^, though a nonlinear reanalysis of this dataset did detect a normalization-mediated negative context effect^34^. Other studies have found positive distracter effects^4,5^, though the robustness of these results has also been questioned^63^ and may vary across individuals^36^. Furthermore, both negative and positive context effects may coexist in the same individual choice dataset^4,5^. This controversy points to a critical gap in our understanding of the underlying neural mechanisms of decision-making.

The significance of the current study lies in its enhancement of the representation of noise in divisive normalization class models as a tool for reconciling these two narratives. By differentiating the sources of noise in the model, our work offers a comprehensive framework that reconciles the conflicting evidence in the field. We demonstrate that the timing of noise—whether it occurs before or after the normalization process—can significantly alter the influence of contextual options. Our findings suggest that a contextual input can enhance decision accuracy under early noise, whereas impairs it when under late noise. This systematic understanding not only clarifies the discrepancies in the literature but also broadens our understanding of neural computations, emphasizing the temporal structure of noise in neural processing and decision-making. The current study, therefore, provides a coherent and physiologically informed explanation of how decisions are shaped by context. While our results show that different patterns of observed context effects^4,5,9,32,34–36^ can be driven by the balance of early and late noise, we note this does not rule out other coexisting mechanisms for varying context effects^5,35,36^. For future research, the current study offers a valuable perspective on examining contextual impacts by isolating the role of contextual variance from contextual mean, a factor often overlooked in the literature.

Before addressing its neurobiological implications, it is important to recognize the limitations of the current experimental design. The current design does not allow us to measure early and late noise during choices directly. Hence, it is worth clarifying our assumptions regarding the underpinning neural computations. The early noise associated with single items was manipulated by controlling the uncertainty of information provided to participants about choice objects. The impact of this manipulation was measured by assessing the variability of bidding behavior. This measured variability was incorporated into the normalization computation as distributions on inputs to the model. The model comparison results support our hypothesis that the itemized variability is processed by normalization (*Model 4*) instead of a linear add-on to the choice stochasticity (*Model 2*). This aligns with a neurobiological mechanism recently uncovered in the study of value-based learning, where individual neurons encode a spectrum of values, shaping the overall signal distributions^64^. A second limitation that must be acknowledged is our assumption that time pressure solely affects the late noise arising from the output of the normalization process. While longer decision times have been suggested to reduce the uncertainty of value representation^65,66^, an additional model we tested by allowing time pressure to affect the itemized variability before normalization performed worse than *Model 4* (*Model 5* vs. *Model 4*: ΔAIC = 33; ΔBIC = 243). It suggests that time pressure affects late noise only instead of both early and late noise.

From an evolutionary perspective, uncertainty in the representation of value poses significant challenges for humans and animals in making accurate decisions. The uncertainty of an option’s value is an inevitable aspect of decision-making and is due to various factors like the stochasticity inherited from the environment^37^, uncertain reward associations^67–70^, stochastic memory retrieval^39,71^, imprecise inferential reasoning^72,73^, and/or fluctuating motivational states^74,75^. Our findings regarding early noise illustrate a special case where noise can be effectively managed by the brain circuit. Divisive normalization emerges as a crucial mechanism that enhances value representations by squeezing the uncertainty of input values with contextual information. In contrast to a more traditional perspective, which focuses on the detrimental effects of context under late noise, the early noise scenario reveals another biological benefit of divisive normalization in real-world environments. Together with other well-documented benefits of divisive normalization, such as range adaptation^21,24,76^ and redundancy elimination^18,22^, the current work highlights the biological value of divisive normalization under noise.

Another implication of the current work is the potential importance of noise correlation. To predict the positive contextual effect observed under early noise, a certain degree of independence between the denominators of different options is required. This implies a biological substrate where inhibitory neurons, a biological counterpart of the divisive gain-control from the denominator^55,56,76^, are distinct pools for different options. This motif is supported by recent discoveries of inhibitory neurons with choice-selective properties^77^ and implemented in models like the local disinhibition decision model (LDDM)^56^. Similarly, in the late noise scenario, when the late noise covariance is fully correlated across options, the choice would lose its stochastic nature. Adding a constant value to all inputs would always preserve their rank during choice, inevitably resulting in 100% choice accuracy, which obviously contradicts empirical observations. Simultaneous recording of multiple units from the visual cortex shows that the trial-by-trial covariance across individual neurons decreases when animals are more engaged in a task and the performance in discriminating between stimuli is enhanced^78^. This implies that the noise structure imposed on the task-relevant signal is inherently decorrelated, aligning with the independent noise in our model.

In conclusion, our extension of the divisive normalization model to incorporate input stochasticity provides new insights into contextual effects in decision-making. This extended model, validated through our behavioral data, reconciles previously conflicting findings that challenged the traditional divisive normalization model. More broadly, this study underscores the importance of considering the temporal structure of noise in cognitive modeling, paving the way for more precise and predictive frameworks in neuroscience.

## Methods

### Participants

Sixty adult participants were recruited through an online platform in the New York area (https://newyork.craigslist.org/). Among these, five participants consistently bid zero for most items, not allowing effective construction of their choice set, thus excluded from the analysis. Fifty-five participants remained after exclusion (29 females, determined based on self-report, age = 37.9±12.5). The sample size was determined by power analysis, which showed that for a medium effect size (Cohen’s D = .5), more than fifty-four participants would be required to detect the difference of context-modulation slopes between conditions within-subject (paired t-test, two-tailed) at a power level of 95% and significance level of .05 (G*Power 3.1)^79^. The study protocol was approved by the Institutional Review Board (IRB) of New York University Grossman School of Medicine. All participants were informed of the potential risk of the task and provided written consent. All participants were compensated with $90 or equivalent depending on their task choices.

### Procedures

#### Overall task structure

The participants performed a variant of the two-stage valuation and trinary choice task previously used to examine context-dependent preferences ^9,32^. In this task, participants first provide their subjective valuations of different consumer items in a bidding-based valuation task. Subsequently, they select their preferred item from trinary choice sets; these choice sets were constructed based on their valuations to quantify the degree of context-dependent choice behavior. Experimental details are provided below.

#### Early noise manipulation

Upon the participants’ arrival, we introduced two levels of representational vagueness (precise or vague) of the consumer goods by controlling the information provided to the participants. The participants were provided with a selection of 36 goods exclusively obtained from a prominent online retail store with a wide range of market values (from $.47 to $99.99) based in the New York area, United States. Half of the items were designated as “precise items” and presented as physical products arranged on a table. Participants were instructed to examine and interact with these items thoroughly for a minimum of 10 minutes and informed that the subsequent task would involve the appearance of a cartoon image corresponding to each observed item. The other half of the items were designated as “vague items,” and were only presented as cartoon images in the task; participants were required to make their selections based on their best estimations derived from their daily life experiences. We expected the participants to have a relatively more precise representation of the precise items than the vague items. The assignment of which half of the items were precise was counterbalanced across the participants. The market values and the categories of the goods between the two halves were carefully matched.

#### Bid task to access subjective values and variability

Subsequently, the participants played an incentive-compatible Becker-Degroot-Marschak (BDM) auction for each good^80^, which was designed to reveal the subjective value for each item as well as the variability of that value. Each participant was provided an initial endowment of $90. In each bidding trial, a cartoon image of an item was presented in the center of the screen. The participants were free to bid on the item continuously from $0 to $90 by moving their computer mouse (**Figure 3c**). They were informed that a random trial would be drawn at the end of the experiment. If they bid higher in that trial than a random number from $0 to $90 (uniformly distributed), they will get the good at that random price; otherwise, they will keep the full amount of endowment but have no chance to get the good. The goods were presented in randomized order, and each item was presented three times in total (108 trials). To quantify the variability of subjective valuations, we calculated the variance across the three bids for each item. In addition, at the end of each trial, participants were asked to rate “how certain you are about the value you bid” on a Likert scale from 0 to 10; certainty ratings showed a high correlation with the bidding variance but are not further analyzed here. Most of the participants completed the bid task in about 10 minutes.

#### Choice task and late noise manipulation

Following the previous tasks, the participants completed an incentive-compatible choice task to measure their choice accuracy between two target options under the influence of a distracter option. In each trial, three goods were presented in counterbalanced locations on the screen (**Figure 3e**). Among the three, two target options were selected from the six top-ranked precise items based on their bid mean values, resulting in fifteen different combinations of target pairs. A third option was selected from the lower-ranked goods, consisting of twelve precise items and another twelve vague items (**Figure 3d**). To manipulate late noise, two levels of time pressure (10 and 1.5 seconds) were implemented in the choice task, alternating in a blockwise manner (block order counterbalanced across participants). Participants were informed at the start of each block about the time limit; failure to make a choice within the time limit would lose the chance of receiving the item. The mean bid values of the distracters were matched between the two time-pressure conditions.

These conditions produced a 2-by-2 design with two levels of early noise (precise and vague items) and two levels of late noise (low and high time pressure), with six distracter items under each condition. Each distracter was combined with the same set of fifteen target pairs across randomized trials, resulting in a total of 90 trials for each condition (360 trials in total). Participants were informed that the choice-task trials would be pooled with the bid-task trials at the end of the experiment, and a random trial from the mixed pool would be drawn for realization. If they received a choice-task trial, they would receive their selected item in that trial; otherwise, if they received a bid-task trial, they would participate in the BDM auction as per the specified rules outlined above.

### Numerical simulations

Simulations were conducted in Matlab R2023a (The Mathworks Inc., 2023). To determine the choice accuracy for each set of option values, 4,096,000 trials were simulated. In each trial, the noise term for each option was randomly drawn from an independently and identically distributed (i.i.d.) *Gaussian* distribution and added either to (1) the option value before divisive normalization (early noise), (2) the product of normalization (late noise), or (3) both (mixed noise), depending on the tests specified in the main text.

It is important to note that the independence of the denominators across options is required to see the positive context effect under early noise. To achieve that, we resampled the value of each option in the denominator independently from their value in the numerator; otherwise, all options would share a fully dependent denominator, resulting in a flat context effect under early noise.

Choice accuracy between the two targets was computed as the ratio of the number of trials when *V_1_* appeared as the maximum value versus the total number of trials when *V_1_* or *V_2_* appeared as the maximum; trials when *V_3_* appeared as the maximum were excluded.

The non-negative biological constraint on each option will lead to different predictions from the situation when allowing the option value with noise to be negative (see **Figures 1d-f** and **2a**). To implement the constraint, we forced each option’s value from both the intermediate stage of normalization with early noise and the final product added with late noise to be non-negative.

The target values in the simulations were set arbitrarily as *V_1_* = 88, *V_2_* = 83; and *V_3_* varied from 0 to *V_1_* given the specific setting of each test required. The maximum response value (*K*) was set arbitrarily at 75 Hz. Changing the scale of the inputs and the maximum response will not impact the results as long as their magnitudes are scaled together with those of the early noise (*ε*) and late noise (*η*), which we chose to match roughly the overall choice accuracy of our participants. Baseline normalization was assumed at a low level of *σ*_*H*_ = 1. The weight of the normalization was assumed at *w* = 1. Parameters that were specific to each figure were listed below.

- **Figures 1b,c**: Four examples of *V_3_* were set in panels valued at 0, 47.4, 71.1, and 79. In the early noise example, *ε* was drawn from *N*(0, 2.8^2^) for every option; in the late noise example, *η* was drawn from *N*(0, 1^2^) for every option.
- **Figures 1d-f**: *V_3_* was sampled from 0 to *V_1_* for 50 equally spaced values. In the early noise example, *ε* ∼ *N*(0, 9^2^) for every option, and *η* was set as zero; in the late noise example, *ε* was set as zero, and *η* ∼ *N*(0, 4^2^) for every option. The scales of early and late noise were picked differently since they reflect the variance of different sources. We picked these values by considering a comparable degree of choice stochasticity they will induce.
- **Figure 2a**: *V_3_* was sampled between 0 and *V_1_* for 50 equally spaced values. The standard deviation of early noise for all options (*σ*_*ε*_) increased from 0 to 9; meanwhile, the standard deviation of late noise for all options ( *σ*_*η*_ ) decreased from 3.63 to 0. The amplitudes of the early and late noise of the lines in the middle were determined by controlling their sum having a constant Euclidean norm.
- **Figures 2b,c**: A wider range of *σ*_*ε*_ and *σ*_*η*_ were tested. *σ*_*ε*_ ranged between 0 and 13, with 101 equally spaced values; *σ*_*η*_ ranged between 0 and 5.24, with 100 equally spaced values. Other parameters kept the same to the above in Figure 2a.
- **Figure 4a-d**: *V_3_* was sampled between 0 and *V_2_* for 50 equally spaced values. The standard deviation of *V_3_*’s early noise varied from 0 to .35\**V_2_*, with six values equally spaced. The early noise of *V_1_* and *V_2_* were fixed at 4.5 to match the overall choice accuracy observed from the data. For the small and large late noise, *σ*_*η*_ for all options were set as 1.0 and 1.4286 to match the effect of time pressure observed from the data.

### Model fitting

Four nested models were compared, varied in their assumptions of the origins of noise and/or whether containing divisive normalization. The simplest model is a probit model (*Model 1*), which assumes a linear coding of each item’s value added with late noise that is indifferently superimposed as a zero-mean *Gaussian* term on every item. The utility of each item leading to probabilistic choice was written in Equation 2,

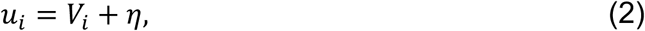

where *V*_*i*_ is the participant’s mean bid on item *i*, 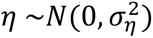 represents the late noise applied identically and independently to all items.

The choice probability of each option out of the three alternatives is determined by a numerical simulation approach, which chooses the option with the maximum value in a single sampling and calculates the chosen ratio of the option over 20,000 times of identical and independent sampling. Negative values were forced to be zero considering the non-negative biological constraint. When multiple options (*n* ∈ [2,3]) occur with the same value that is a maximum, these options were counted with an equal vote of 1/*n*, although these were rare cases in our simulations.

In fitting the chosen ratio, Equation 2 is equivalent to fit Equation 3 by standardizing the deviance of noise and scaling the item’s value by 1/*σ*_*η*_,

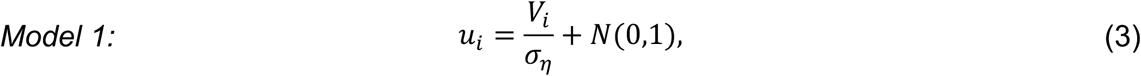

An extension of the assumption on the noise is to incorporate early noise, reflecting the representational uncertainty of each item; while value coding stays linear (Equation 4) (*Model 2*),

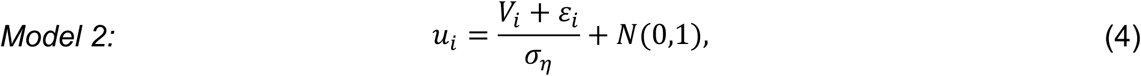

Different from the late noise *η* that is indifferently applied to all items, the early noise 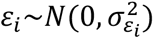 is specific to every single item. In our dataset, we used the measured bidding variance on every item to approximate the early-noise variance, with a scaling parameter *λ* fitted individually (Equation 5),

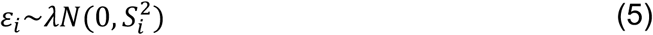

where *S*_*i*_ is the measured bidding standard deviation of item *i*. *Model 2* (Equation 4) can easily collapse to *Model 1* (Equation 3) when *λ* = 0.

Other than the early noise, we incorporated divisive normalization as a potential non-linear mechanism of value coding (Equation 6),

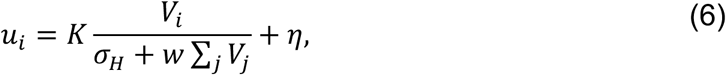

Similar to Equation 1 in the main text, the option’s value is normalized by a weighted sum of all options in the choice set and a constant scaling parameter *σ*_*H*_. Equation 6 is equivalent to Equation 7 by standardizing the late noise and absorbing the constant scale to the denominator (*Model 3*),

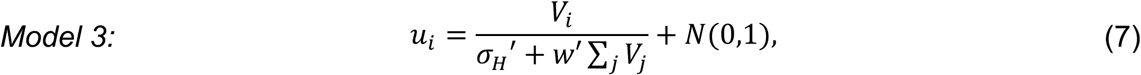

where *σ*_*H*_′ is a composite index reflecting the value signal relative to the late noise; *w*′ captures any potential context effects regardless of the scaling. *Model 3* (Equation 7) easily degrades to *Model 1* (Equation 3) when *w*^′^ = 0.

For our dual-noise divisive normalization model, we incorporated early and late noise into divisive normalization (Equation 8) (*Model 4*),

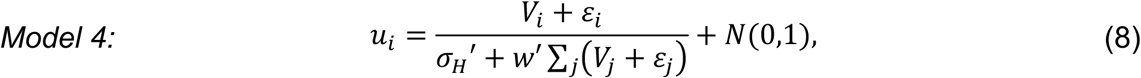

where the early noise term *ε*_*i*_ shares the same setting as specified in Equation 5. *Model 4* easily degrades to *Model 3* when *λ* = 0 and *w*^′^ > 0, to *Model 2* when *w*^′^ = 0 and *λ* > 0, and to *Model 1* when *w*^′^ = 0 and *λ* = 0.

Considering that early noise is specific to each item and significantly different between the vague and precise items, we did not use additional parameters to capture the early noise difference between the vague and precise conditions. While, late noise, which is expected to be shaped by time pressure, would have a systematic difference between the low and high time-pressure conditions. To capture that, we utilized a dummy variable *δ* to set the low time-pressure condition as a baseline and fit the potential difference between the high time-pressure condition (Equation 9),

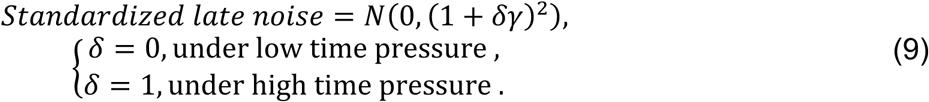

where *γ* captures the potential difference of late noise magnitude in the high time-pressure condition to the low time-pressure condition.

With that, we have 2 parameters for Model 1 (*σ*_*η*_, *γ*), 3 parameters for *Model 2* (*σ*_*η*_, *γ*, *λ*) and *Model 3* (*σ*_*H*_′, *γ*, *w*′), and four parameters for Model 4 (*σ*^′^, *γ*, *w*^′^, *λ*). We fitted those models individually with maximum likelihood and optimized with Bayesian Adaptive Direct Search (BADS) algorithm in Matlab^81^. The BADS algorithm utilizes the features of plausible range and hard boundary for each parameter to optimize its performance of estimation. We set these values by roughly estimating their impact on choice accuracy and taking multiple times of model fitting attempts, and set the plausible range roughly as the 5th and 95th percentiles of each parameter and the hard boundary far off the distribution to prevent the parameters from hitting the boundary indicated in Table 2.

The parameters in the denominators (*σ*_*η*_ *or σ*_*H*_′ and *w*′) were restricted to be non-negative to prevent negative utility values. The scaling parameter on early noise (*λ*) was restricted to be non-negative since variance cannot be negative.

In addition, to test whether time pressure could affect the contribution of early noise to choice accuracy other than on late noise, we allowed time pressure to affect the itemized variance before normalization, as indicated in Equation 10 (*Model 5*). Comparison between *Model 5* and *Model 4* revealed that *Model 5* performed worse than *Model 4* in capturing participants’ choices (*Model 5* vs. *Model 4*: ΔAIC = 33; ΔBIC = 243), suggesting that time pressure affects late noise only rather than both early and late noise.

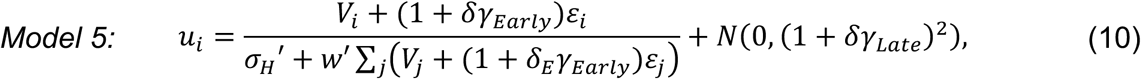

### Post-hoc test on the linear trends

Linear regression models were performed on the target-chosen trials pooled across subjects to reveal the linear trends of contextual modulation with the “glm” function in *R* (Version 4.3.1)^82^ by setting the family as binomial. The standardized coefficients and their 95% confidence intervals of corresponding effects reported were achieved by using the *R* package “effectsize” (version 1.0.0)^83^.

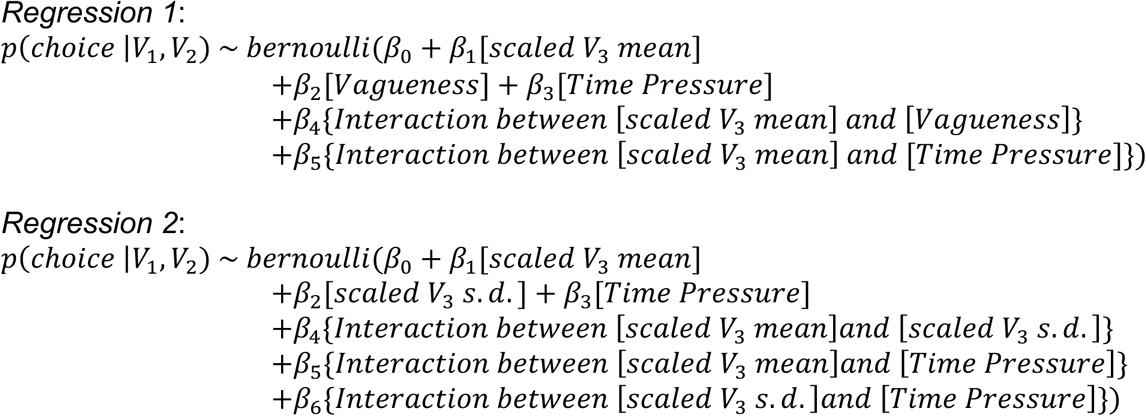

## Data availability

The raw behavioral data generated from this study is available with public access in the GitHub repository https://doi.org/10.5281/zenodo.14940477.

## Code availability

The code for performing simulations and generating figures in this study is available with public access in the GitHub repository: https://doi.org/10.5281/zenodo.14940477.

## Acknowledgments

The authors thank Dr. Ryan Webb, Miss Aliya Koishina for their constructive comments and feedback. This study is supported by NIH U19 NS107616 (to K.L.).

## Author contributions

B.S., P.W.G., and K.L. conceptualized the idea. B.S. and J.W. designed the experiment. B.S., J.W., and D.N. performed the experimental studies. B. S. carried out the data analysis and the computational studies. B.S. and D.N. carried out the analytical proof. P.W.G. and K.L. co-supervised the work.

