## Supplementary Information for "Early versus late noise differentially enhances or degrades context-dependent choice"

### *Supplementary Note 1*

This appendix provides an analytical analysis of the generality of the effects we show in the simulations of the main text. Since divisive normalization raises a challenging mathematical issue regarding the *Ratio of Gaussians* distribution[1–3], we took an approximation to the *Ratio of Gaussians* in our analysis using the method reported in the literature[4] to provide a qualitative understanding of the generality of the results. Future work will be required to derive a closed-form solution using the *Ratio of Gaussians* approach, if such a solution exists.

In this analysis, we assumed that both early and late noise are represented as zero-mean *Gaussian* distributions, independent of each other since they have independent biological origins. Additionally, each noise term is assumed to be independent across options. When the neural activity of every option is much higher than zero, the non-negative constraint on its distribution is neglectable, and the represented value can thus be written as,

$$FR_i = \frac{V_i + \varepsilon_{i,n}}{\sigma_H + \sum_j w(V_j + \varepsilon_{i,d})} + \eta_i, \qquad (1)$$

where $\varepsilon_{i,n} \sim N\left(0, \sigma_{\varepsilon_i}^2\right)$ is the early noise of option $i$ $(i \in [1, \ldots, N], N \geq 2)$ in the numerator, $\varepsilon_{i,d} \sim N\left(0, \sigma_{\varepsilon_i}^2\right)$ is the early noise of option $i$ in the denominator; they are marked differently considering they may have covariance (i.e., they are represented by two different populations of neurons: $Cov\left(\varepsilon_{i,n}, \varepsilon_{i,d}\right) = \rho\sigma_{\varepsilon_i}^2$); $\eta_i \sim N\left(0, \sigma_{\eta_i}^2\right)$ is the late noise of option $i$ and assumed to be independent of early noise. The covariance structure of all noise terms across options is summarized in Supplementary Equation 2, with most of the terms independent from each other. Nonzero across-option covariance terms will significantly increase the complexity of the analysis, and we assume it to be zero here; we acknowledge this is an important topic to discover in the future about the covariance structure of noise in population coding.

$$\begin{pmatrix} \varepsilon_{i,n} \\ \varepsilon_{i,d} \\ \eta_i \\ \vdots \\ \varepsilon_{j,n} \\ \varepsilon_{j,d} \\ \eta_j \end{pmatrix} \sim N\left( \begin{pmatrix} 0 \\ 0 \\ 0 \\ \vdots \\ 0 \\ 0 \\ 0 \end{pmatrix}, \begin{pmatrix} \sigma_{\varepsilon_i}^2 & \rho\sigma_{\varepsilon_i}^2 & 0 & & 0 & 0 & 0 \\ \rho\sigma_{\varepsilon_i}^2 & \sigma_{\varepsilon_i}^2 & 0 & \cdots & 0 & 0 & 0 \\ 0 & 0 & \sigma_{\eta_i}^2 & & 0 & 0 & 0 \\ & \vdots & & \ddots & & \vdots & \\ 0 & 0 & 0 & & \sigma_{\varepsilon_j}^2 & \rho\sigma_{\varepsilon_j}^2 & 0 \\ 0 & 0 & 0 & \cdots & \rho\sigma_{\varepsilon_j}^2 & \sigma_{\square\square_j}^2 & 0 \\ 0 & 0 & 0 & & 0 & 0 & \sigma_{\eta_j}^2 \end{pmatrix} \right) \tag{2}$$

We defined the terms from Supplementary Equations 1 and 2 as,

$$FR_i = \frac{N_i}{D_i} + L_i, \tag{3}$$

where $N_i \sim N(V_i, \sigma_{\varepsilon_i}^2)$ is the direct input with early noise in the numerator, $D_i \sim N\left(S, \sigma_{D_i}^2\right)$ is the denominator integrating all options, and $L_i$ indicates the additive zero-mean Gaussian distribution for the late noise. Contained in the denominator, $S = \sigma_H + w \sum_i V_i$ is defined as the mean value of the denominator remaining constant across options. $\sigma_{D_i}^2 = w^2\left(\sigma_{\varepsilon_1}^2 + \sigma_{\varepsilon_2}^2 + \sigma_{\varepsilon_3}^2\right)$ indicates the variance of the denominator when considering the independence of options. The covariance between the numerator and denominator is noted as $Cov(N_i, D_i) = \rho w \sigma_{\varepsilon_i}^2$, considering that there may have covariance between the two populations of neurons representing the excitability of the numerator and providing inhibition in the denominator.

#### Under early noise

With only early noise, the representation of each option follows a distribution of the *Ratio of Gaussians* (Supplementary Equation 4),

$$FR_i \sim \frac{N(V_i, \sigma_{\varepsilon_i}^2)}{N\left(S, \sigma_{D_i}^2\right)}, \tag{4}$$

Here we took an approximation depending on Taylor expression around $(V_i, S)$ and neglecting the higher-order terms [4]. This approach allows us to approximate the *Ratio of Gaussians* with a *Gaussian* distribution (Supplementary Equation 5),

$$FR_i \simeq N(\mu_i, \sigma_i^2), \tag{5}$$

where $\mu_i \simeq \frac{V_i}{S}$, and $\sigma_i^2 \simeq \frac{V_i^2}{S^2}\left(\frac{\sigma_{\varepsilon_i}^2}{V_i^2} + \frac{\sigma_{D_i}^2}{S^2} - 2\rho w \frac{\sigma_{\varepsilon_i}^2}{V_i S}\right)$.

Thus, the signal-to-noise ratio of each single item, marked as $SNR_i$, can be approximated as Supplementary Equation 6,

$$\begin{aligned} SNR_i &\cong \frac{\mu_i}{\sigma_i} \\ &= \left(\frac{\sigma_{\varepsilon_i}^2}{V_i^2} + \frac{\sigma_{D_i}^2}{S^2} - 2\rho w \frac{\sigma_{\varepsilon_i}^2}{V_i S}\right)^{-\frac{1}{2}} \end{aligned} \tag{6}$$

Supplementary Equation 6 can be proved to be always non-negative, regardless of the values of $\rho$ and $w$.

We examined how $SNR_i$ changes with a contextual mean value $V_j$ by examining the partial derivative (Supplementary Equation 7),

$$\frac{\partial \frac{\mu_i}{\sigma_i}}{\partial V_j} = \frac{w}{S^2}\left(\frac{\sigma_{\varepsilon_i}^2}{V_i^2} + \frac{\sigma_{D_i}^2}{S^2} - 2\rho w \frac{\sigma_{\varepsilon_i}^2}{V_i S}\right)^{-\frac{3}{2}}\left(\frac{\sigma_{D_i}^2}{S} - \rho w \frac{\sigma_{\varepsilon_i}^2}{V_i}\right) \tag{7}$$

The sign of the partial derivative is determined by the sign of $\frac{\sigma_{D_i}^2}{S} - \rho w \frac{\sigma_{\varepsilon_i}^2}{V_i}$, which requires the denominator to have a larger fano factor than the numerator to improve the target representation with the contextual mean when $\rho = 1$. However, reducing the numerator-denominator covariance, i.e., when $\rho < 1$, releases such a restriction. When $\rho = 0$, the contextual mean always improves the target representation.

We examined how $SNR_i$ changes with a contextual variance $\sigma_{\varepsilon_j}$ by using the same approach (Supplementary Equation 8),

$$\frac{\partial \frac{\mu_i}{\sigma_i}}{\partial \sigma_{\varepsilon_j}} = -\frac{w}{S^2}\left(\frac{\sigma_{\varepsilon_i}^2}{V_i^2} + \frac{\sigma_{D_i}^2}{S^2} - 2\rho w \frac{\sigma_{\varepsilon_i}^2}{V_i S}\right)^{-\frac{3}{2}} \sigma_{D_i} \tag{8}$$

Supplementary Equation 8 stays negative when $w > 0$ and early noise is above zero. Thus, increasing the contextual noise will always decrease the *SNR* of the individual item(s) under early noise.

To study the choice accuracy between the targets under the three-option set, we examined the discriminability of the two targets $FR_1$ and $FR_2$ using the $d'$ index,

$$d' = \frac{|\mu_1 - \mu_2|}{\sqrt{\sigma_1^2 + \sigma_2^2}} \tag{9}$$

We examined how $d'$ changes with a contextual mean value $V_3$ by examining the partial derivative (Supplementary Equation 10),

$$\frac{\partial d'}{\partial V_3} = \frac{w}{S^3} \frac{|V_1 - V_2|}{(\sigma_1^2 + \sigma_2^2)^{\frac{3}{2}}} \Big( (\sigma_{\varepsilon_1}^2 + \sigma_{\varepsilon_2}^2 + \sigma_{\varepsilon_3}^2)(V_1^2 + V_2^2) - \rho w S \big( V_1 \sigma_{\varepsilon_1}^2 + V_2 \sigma_{\varepsilon_2}^2 \big) \Big) \tag{10}$$

The sign of the partial derivative is determined by $(\sigma_{\varepsilon_1}^2 + \sigma_{\varepsilon_2}^2 + \sigma_{\varepsilon_3}^2)(V_1^2 + V_2^2) - \rho w S\big(V_1 \sigma_{\varepsilon_1}^2 + V_2 \sigma_{\varepsilon_2}^2\big)$, which is positive when $\rho = 0$ and can be either positive or negative when $\rho > 0$.

We examined how $d'$ changes with a contextual variance $\sigma_{\varepsilon_3}$ by examining the partial derivative (Supplementary Equation 11),

$$\frac{\partial d'}{\partial \sigma_{\varepsilon_3}} = -\frac{w^2}{S^2} \frac{|V_1 - V_2|}{(\sigma_1^2 + \sigma_2^2)^{\frac{3}{2}}} (V_1^2 + V_2^2) \sqrt{\sigma_{\varepsilon_1}^2 + \sigma_{\varepsilon_2}^2 + \sigma_{\varepsilon_3}^2} \tag{11}$$

The sign of partial derivative stays negative as long as $w > 0$ and the targets have mean values and early noise above zero.

**Remarks**

1. According to Supplementary Equation 7, when the early noise terms of an option in the numerator and in the denominator are independent, i.e., $Cov\big(\varepsilon_{i,n}, \varepsilon_{i,d}\big) = 0$ , the representation of an individual item under contextual modulation (when $w > 0$) will be improved as the distracter value increases. In other words, the neural representation of the value of a given option will show a higher *SNR* with increasing contextual mean ($V_j$) given a fixed contextual variance ($\sigma_{\varepsilon_j}$). This is what we observed in the numerical simulations of the main text. During the implementation, the early noise terms in the numerator and denominator were independent draws from the generating distribution. In this case, as shown in the analyses above, distracter value will always show a positive context effect under early noise. If these early noise terms are not independent (the numerator-denominator covariance is non-zero), this facilitation effect is not guaranteed and will depend on the ratios of fano factors between the numerator and the denominator.

2. According to Supplementary Equation 8, when $w > 0$ (i.e. when divisive normalization occurs), the representation of a target item will always be impaired under larger contextual variance, exhibiting a lower *SNR*.

3. According to Supplementary Equation 10, when $w > 0$, the discriminability of the two targets will always be facilitated by an increasing mean of the third option if the early noise terms in the numerator and the denominator are independent (i.e., $\rho = 0$). This was the assumption implemented in the simulations in the main text. Note that this facilitation effect is not guaranteed if $\rho > 0$ and distracter value can either facilitate or impair target choice, as indicated in Supplementary Equation 10. Thus, independence of the early noise terms in the numerator and denominator is critical to predict contextual facilitation.

4. According to Supplementary Equation 11, when $w > 0$, the discriminability of the two targets will always be impaired under larger contextual noise, consistent with what we observed in our empirical data.

**Under late noise**

With only late noise, the representation of each option follows a *Gaussian* distribution (Supplementary Equation 12),

$$FR_i \sim N\left(\frac{V_i}{S}, \sigma_{\eta_i}\right), \tag{12}$$

The $SNR_i$ decreases with a contextual mean value $V_j$ since the partial derivative (Supplementary Equation 13) is always negative when $w > 0$.

$$\frac{\partial \frac{V_i}{S\sigma_{\eta_i}}}{\partial V_j} = -\frac{w}{S^2}\frac{V_i}{\sigma_{\eta_i}} \tag{13}$$

The $SNR_i$ will not be impacted by the contextual late noise since the late noise terms are fully independent and not involved in normalization (Supplementary Equation 14).

$$\frac{\partial \frac{V_i}{S\sigma_{\eta_i}}}{\partial \sigma_{\eta_j}} = 0 \tag{14}$$

By checking the discriminability of the two targets in the special case of three options, the $d'$ index always decreases with the contextual value $V_3$ when $w > 0$ (Supplementary Equation 15).

$$\frac{\partial d'}{\partial V_3} = -\frac{w}{S^2}\frac{|V_1 - V_2|}{\sqrt{\sigma_{\eta_1}^2 + \sigma_{\eta_2}^2}} \tag{15}$$

The discriminability of the two targets will not be impacted by the late noise of the third option (Supplementary Equation 16).

$$\frac{\partial d'}{\partial \sigma_{\eta_3}} = 0 \tag{16}$$

**Remarks**

1. Under late noise, when $w > 0$, increasing the mean value of the third option will always impair the representation of each single target as well as the discriminability between the two targets. This aligns with the standard view of negative context effects presented in the literature[5].

2. The variance of the third option under a late noise assumption will have no impact on the targets, in contrast to the evidence in the main text, suggesting the involvement of a process other than late noise.

**Under mixed noise**

With both early and late noise, the representation of each option can be approximated as the combination of the *Ratio of Gaussians* term contributed by early noise with another zero-mean *Gaussian* term contributed by late noise (Supplementary Equation 17),

$$FR_i \simeq N\left(\mu_i, \sigma_i^2 + {\sigma_\eta}_i^2\right), \tag{17}$$

where $\mu_i \simeq \frac{V_i}{S}$ and $\sigma_i^2 \simeq \frac{V_i^2}{S^2}\left(\frac{{\sigma_\varepsilon}_i^2}{V_i^2} + \frac{\sigma_{D_i}^2}{S^2} - 2\rho w \frac{\sigma_{\varepsilon_i}^2}{V_i S}\right)$.

The $SNR_i$ of a single item can be approximated as in Supplementary Equation 18,

$$\frac{\mu_i}{\sqrt{\sigma_i^2 + \sigma_{\eta_i}^2}} = \left(\frac{{\sigma_\varepsilon}_i^2}{V_i^2} + \frac{\sigma_{D_i}^2}{S^2} - 2\rho w \frac{\sigma_{\varepsilon_i}^2}{V_i S} + \frac{S^2}{V_i^2}\sigma_{\eta_i}^2\right)^{-\frac{1}{2}} \tag{18}$$

With the additional term in Supplementary Equation 18 contributed by late noise (compared to Supplementary Equation 6), the partial derivative of *SNR* with respect to contextual mean value $V_j$ contains an additional term (Supplementary Equation 19),

$$\frac{\partial \frac{\mu_i}{\sqrt{\sigma_i^2 + \sigma_{\eta_i}^2}}}{\partial V_j} = \frac{w}{S^2}\left(\frac{{\sigma_{\varepsilon_i}}^2}{V_i^2} + \frac{\sigma_{D_i}^2}{S^2} - 2\rho w \frac{\sigma_{\varepsilon_i}^2}{V_i S} + \frac{S^2}{V_i^2}\sigma_{\eta_i}^2\right)^{-\frac{3}{2}} \cdot \left(\frac{\sigma_{D_i}^2}{S} - \rho w \frac{\sigma_{\varepsilon_i}^2}{V_i} - \frac{S^3}{V_i^2}\sigma_{\eta_i}^2\right) \tag{19}$$

With the additional negative term contributed by late noise, Supplementary Equation 19 shows a tradeoff between early and late noise in driving opposite context effects: the contextual mean under early noise improves $SNR_i. In\ contrast,$ late noise leads to an impairment effect, in addition to the contribution of numerator-denominator covariance we discussed in Supplementary Equation 7.

Taking the derivative of contextual early noise over $SNR_i$ resulted in the same form as Supplementary Equation 8. Thus, increasing contextual early noise will also impair the target representation.

To examine the accuracy of the choice between the targets in three-option choices, the discriminability of the two targets changes with respect to contextual mean value. This shows a tradeoff between early and late noise amplitudes (Supplementary Equation 20): The discriminability of the two targets will always be improved with $V_3$ under early noise when $\rho = 0$, while increasing the late noise will lead to an impairment effect.

$$\frac{\partial d'}{\partial V_3} = \frac{w}{S^3} \frac{|V_1 - V_2|}{\left(\sigma_1^2 + \sigma_{\eta_1}^2 + \sigma_2^2 + \sigma_{\eta_2}^2\right)^{\frac{3}{2}}} \Big(\left(\sigma_{\varepsilon_1}^2 + \sigma_{\varepsilon_2}^2 + \sigma_{\varepsilon_3}^2\right)(V_1^2 + V_2^2) - \rho w S\left(V_1\sigma_{\varepsilon_1}^2 + V_2\sigma_{\varepsilon_2}^2\right) - S^4\left(\sigma_{\eta_1}^2 + \sigma_{\eta_2}^2\right)\Big) \tag{20}$$

**Remarks**

1. With mixed noise, whether increasing the contextual mean value facilitates or impairs target representation will rely on the relative magnitudes of early and late noise. According to Supplementary Equations 19 and 20, larger summed early noise across options and smaller late noise of the target will lead to a more positive context effect on the target representation and choice accuracy between the targets, consistent with our empirical observations.

2. Under mixed noise, increasing the contextual early noise always impairs the target representation and discriminability, which aligns with our empirical observations.

### *Supplementary Note 2*

The main text demonstrated how the distracter mean value impacts the conditional choice accuracy between the targets under constant levels of distracter early noise. However, the participants' bidding behavior showed that distracter variance scales with distracter mean value (**Figure 3f**). To fill this gap, here we provided simulations of contextual effects incorporating mean-scaled variance.

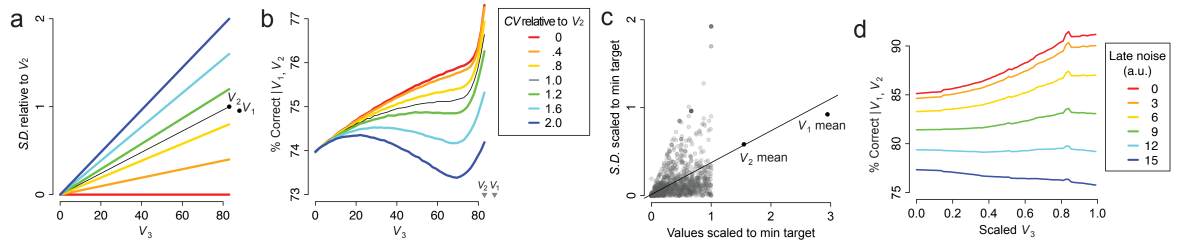

**Supplementary Figure 1. Model predictions under mean-scaled early noise demonstrate positive contextual effects. a** An illustration of the input values and their early noise used in the divisive normalization model without late noise. The distracter (*$V_3$*) ranges from 0 to the lower-valued target *$V_2$*. Six levels of mean-variance ratio were tested, as indicated in the colored lines. The y-axis shows the standard deviation of early noise scaled to the early noise of *$V_2$*. The thin dark line indicates the mean-variance ratio of *$V_2$*. The values and scaled variance of the two targets were indicated by text. **b** Model predictions of conditional choice accuracy between the targets as a function of *$V_3$*, evaluated across six levels of mean-variance ratios. Each curve corresponds to a coefficient of variation (*CV*) of the distracters ($\sigma_{\varepsilon_3}/V_3$) relative to the *CV* of the target ($\sigma_{\varepsilon_2}/V_2$), indicated by the thin dark line. The model predicts positive contextual effects when the *CV* of the distracters is under the level of the target and gradually leads to negative effects when the *CV* of the distracters exceeds the level of the target. **c** Mean values and standard deviations of items from the participants' bidding data. Single distracters from all participants were visualized as a cloud of dots, scaled to the minimum target value within each participant. The mean values and standard deviations of the targets averaged across all participants were indicated by dots with text annotations. The line represents a linear fit of the ratio between the standard deviation and the mean of *$V_3$*. **d** Model predictions incorporating the empirical data structure. The top red curve indicates the model predictions without late noise. While subsequent curves correspond to increasing levels of late noise, with their arbitrary values indicated in the legend.

We tested two sets of parameters for the simulations. The first set of parameters aligns with the parameters used in the simulations of the main text (**Figures 4a-d**), with fixed target values (*$V_1$* = 88, *$V_2$* = 83) and target early noise ($\sigma_{\varepsilon_1}, \sigma_{\varepsilon_2} = 4.5$). Distracter values ranged from 0 to *$V_2$*. We tested six ratios of the distracter's standard deviation to its mean ($\sigma_{\varepsilon_3}/V_3$), from 0 to $2\sigma_{\varepsilon_2}/V_2$. The largest early noise in the distracters was twice the early noise of the target *$V_2$*. **Supplementary Figure 1a** visualized such mean-variance ratios, with the y-axis normalized to $\sigma_{\varepsilon_2}$. Results indicate that when the mean-variance ratio of the distracters is below the level of *$V_2$* (thin dark line in **Supplementary Figure 1b**), the divisive normalization model with early noise predicts positive contextual effects on the conditional choice accuracy (colored lines above the dark line); when the ratio exceeds the target level, contextual effects could lead to non-monotonic pattern and impair choices when $V_3$ and its variance are large (colored lines below the dark line). Under the current study tested conditions, where the target values are always larger than the distracters, distracters' early noise likely remains below the target level. Thus, the model with mean-scaled early noise still predicts positive contextual effects.

The second set of tested parameters incorporated the mean and variance structure from the empirical data, visualized in **Supplementary Figure 1c**. The mean and standard deviation of the distracters, scaled to the minimum target value for each individual, were shown in the cloud of dots with a fitted line representing the mean-variance ratio of the distracters. The average values and variance of the two targets were indicated on the right side of the panel, closely following the mean-variance ratio of the distracters. The predicted choice accuracy from the divisive normalization model was visualized using the same sliding window approach from the main text (**Figure 4e**) in **Supplementary Figure 1d** (the four parameters of *Model 4* in the simulation: $\sigma_H' = 1, w' = 1, \gamma = 0, and\ \lambda = 1$). Without late noise, the model predicts a positive contextual effect (the red curve). By introducing late noise, such a positive effect wanes and leads to a negative contextual effect when late noise increases (graded colors).

### *Supplementary Note 3*

We provide an alternative method to define the vagueness of the distracters by performing a median split on their bidding variance. Linear regression, as defined below in *Regression 3*, revealed a significant decline in choice accuracy with the newly defined label of distracter vagueness ($\beta$ = -.13, s.e. = .05, $t$ = -2.40, $p$ = .016, standardized coefficient = -.04, 95% C.I. = [-.11, .03]). The mean value of scaled $V_3$ showed a significant interaction with this vagueness, indicating that the contextual slope was more positive under vaguer distracters ($\beta$ = .28, s.e. = .12, $t$ = 2.31, $p$ = .021, standardized coefficient = .09, 95% C.I. = [.01, .16]), consistent with the model prediction. Additionally, time pressure led to a significant decline in choice accuracy ($\beta$ = -.17, s.e. = .05, $t$ = -3.50, $p$ < .001, standardized coefficient = -.17, 95% C.I. = [-.71, -.59]).

*Regression 3*:

$$p(choice\ |V_1, V_2) \sim bernoulli(\beta_0 + \beta_1[scaled\ V_3\ mean] \\ +\beta_2[Vagueness\ new\ label] + \beta_3[Time\ Pressure] \\ +\beta_4\{Interaction\ between\ [scaled\ V_3\ mean] and\ [Vagueness\ new\ label]\} \\ +\beta_5\{Interaction\ between\ [scaled\ V_3\ mean] and\ [Time\ Pressure]\})$$

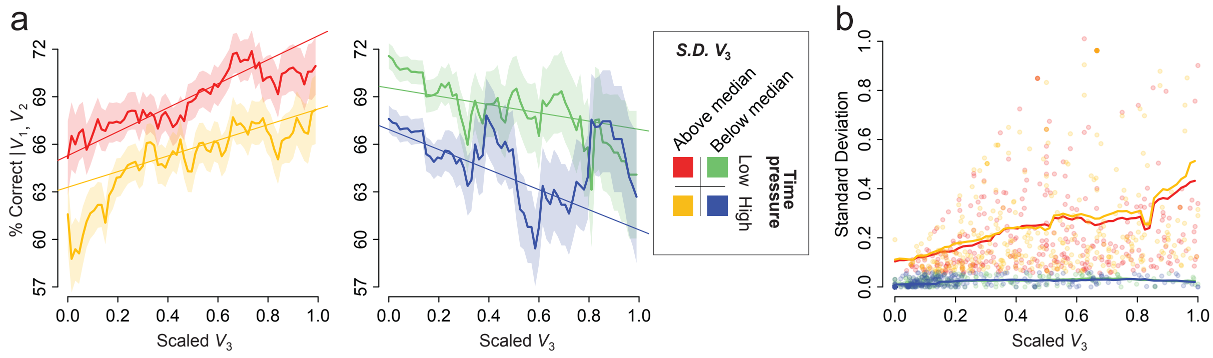

**Supplementary Figure 2. Aggregating the data according to the bidding variance of $V_3$ resulted in a pattern consistent with the experimental conditions of vagueness. a** Pooling the data by a median split of $V_3$'s bidding variance and visualizing the mean choice accuracy (solid lines) using a sliding window approach (window span = .3, step size = .015 over scaled $V_3$; shaded areas indicate the standard deviation of choice probability at each window) revealed a consistent pattern: a facilitation effect when $V_3$'s variance is large and an impairment effect when $V_3$'s variance is small. Linear trends were indicated by the straight lines fitted in the constrained range of scaled $V_3$ from .2 to .8. **b** Visualization of $V_3$'s mean and standard deviation by median split. Each dot represents a $V_3$ item, and the lines indicate the running mean of $V_3$'s standard deviation over the sliding window of scaled $V_3$ (window span = .3, step size = .015).

### *Supplementary Note 4*

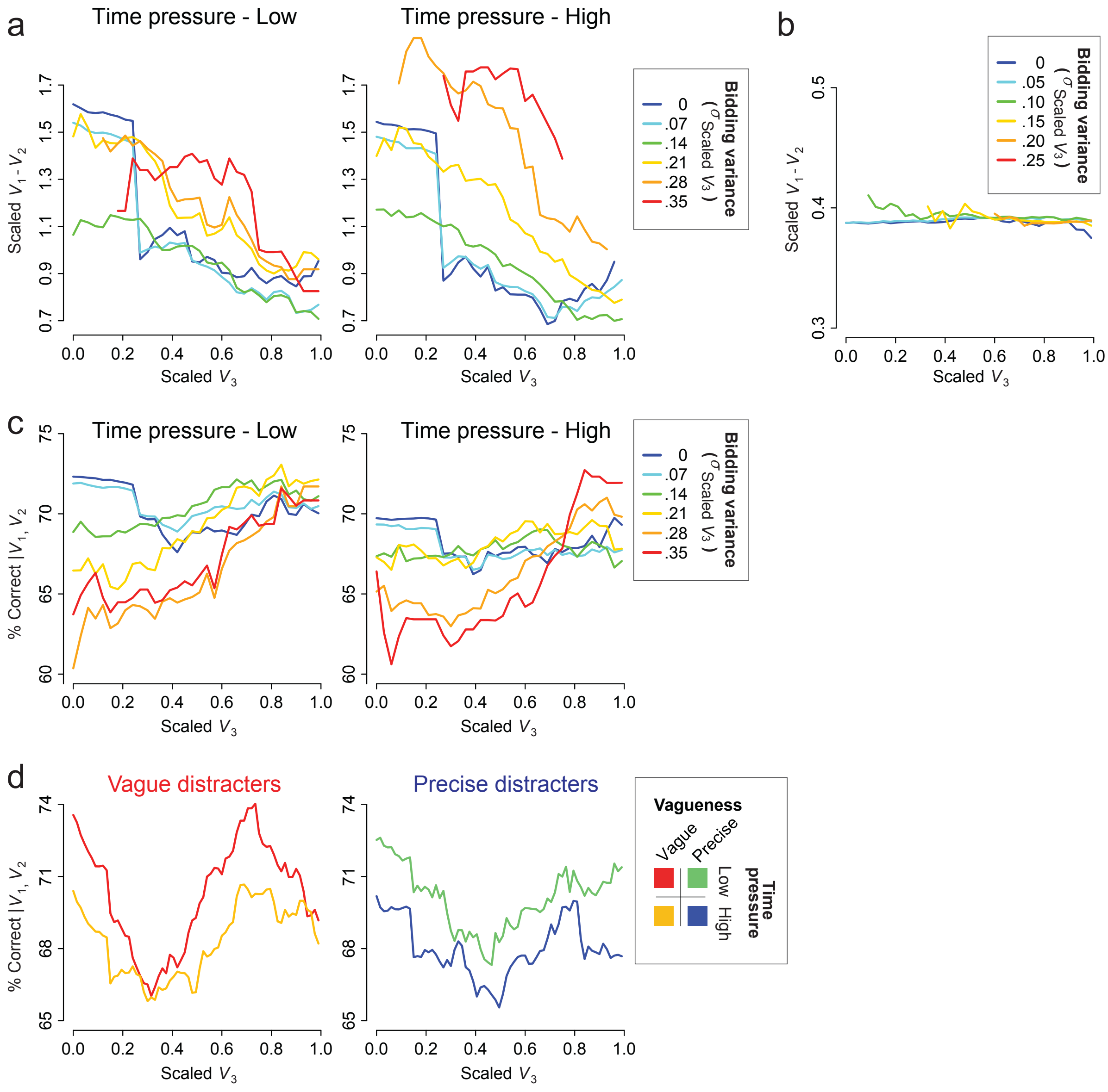

**Supplementary Figure 3. The data structure caused by the pooling process was confirmed as random through simulations and was captured well by the best-fit model. a** The difference between the scaled values of the two targets was aggregated using a sliding-window pooling process based on scaled $V_3$ (sliding window width = .5, step size = .03) and the standard deviation of scaled $V_3$ (sliding window width = .14, step size = .07). This revealed a co-varying structure between the value difference and $V_3$ variance, where larger $V_3$ variance corresponded with a larger value difference between targets, potentially causing a positive shift in choice accuracy for higher $V_3$ variance. **b** Simulations of 5000 subjects, using the same bidding process and mean-scaled noise mechanism, showed no systematic structure in the data, confirming that the positive shift observed in the dataset was caused by random. **c** The posterior predictive check plot from *Model 4*, which fitted the data best among alternative models, showed that the model can capture the positive shift in the data structure, particularly when scaled $V_3$ was large. **d** The same predictive data from the best-fitting *Model 4* but aggregated according to the 2-by-2 experimental conditions. The model, considering the mean and variance structure of the data, captures the aggregated empirical pattern well (see **Figure 4e**).
